# The Red Blood Cell Surface Defines a Distinct Tumor DNA Compartment for Liquid Biopsy

**DOI:** 10.64898/2026.10.07.755430

**Authors:** Elias Elias, Henry Ruger, Jarrod Predina, Benjamin A. Bleiberg, Phurin Areesawangkit, Estela Noguera-Ortega, Abhishek Kulkarni, Jeffrey C. Thompson, Michael G. S. Shashaty, Corey Langer, Steven M. Albelda, Nilam S. Mangalmurti

## Abstract

Red blood cells (RBCs) modulate host inflammatory responses by scavenging mitochondrial and pathogen DNA. However, how RBC surface properties govern the capture of tumor-derived DNA remains poorly understood. We show that RBCs tether and retain tumor-derived DNA on their surface, forming a distinct circulating pool of RBC-bound tumor DNA (rtDNA). Desialylation increased DNA capture in a Toll-like receptor 9 (TLR9)-dependent manner, identifying RBC surface sialylation and TLR9 as regulators of DNA binding. In mice bearing lung cancer xenografts, rtDNA was detected in all animals and its abundance correlated with tumor burden, whereas tumor DNA was undetectable in matched plasma samples. In patients with lung cancer, the majority of detected mutant tumor DNA was associated with RBCs rather than plasma, and RBC-derived mutations were concordant with molecular alterations identified in matched tumor tissue. Collectively, these findings establish the RBC surface as a tunable site of tumor DNA capture and retention and challenge the plasma-centric paradigm of liquid biopsy by identifying a distinct cellular reservoir of tumor-derived genetic material.

## Introduction

Red blood cells (RBCs) have long been defined by their role in oxygen transport and gas exchange. Our previous work expanded this view by establishing noncanonical immune functions of RBCs, including their ability to bind and sequester extracellular nucleic acids(*1–5*). This DNA binding capacity coupled with their abundance and continuous transit through the microvasculature position RBCs to capture genetic material released from tissues throughout the body. Yet, RBC-associated DNA remains a largely unexplored compartment for liquid biopsy.

Current liquid biopsy approaches primarily rely on the detection of circulating tumor DNA (ctDNA) within the plasma fraction of blood. ctDNA consists of fragmented nucleic acids released from apoptotic or necrotic tumor cells into the circulation and has emerged as a promising tool for cancer diagnosis and monitoring(*6, 7*). However, several limitations restrict the clinical sensitivity of ctDNA-based assays including the low abundance of ctDNA in circulation, transient stability, and rapid clearance from plasma(*8*). Several retrospective studies have identified ctDNA assay concordance rates with tissue next-generation sequencing (NGS) ranging from 59.1 to 77.6%(*9–13*), highlighting limitations in the sensitivity of current plasma-based approaches across both early-stage and metastatic disease. As a result, there remains a need for alternative blood-based biomarkers with improved sensitivity and stability that can better inform a range of clinical applications, including early detection of actionable molecular targets, personalized treatment selection, and therapeutic monitoring. Recent studies from our group have identified RBCs as a previously underrecognized reservoir of nucleic acids, including tumor-derived DNA(*1–5*). However, important questions regarding the mechanisms underlying RBC-mediated DNA acquisition, stability, sensitivity, and the degree of inter-individual variability in tumor DNA uptake across RBC populations remain uninvestigated. Defining these parameters is essential for establishing rtDNA as a reproducible and clinically translatable source of tumor-derived genetic information for cancer detection and disease monitoring.

Here, we identify the RBC surface as a tunable and dynamic interface for tumor DNA, which we term RBC-bound tumor DNA (rtDNA). We show that RBCs capture tumor DNA within minutes of exposure and rtDNA could be detected from as few as 1,000 cells, revealing an unexpectedly sensitive nucleic acid reservoir. We further demonstrate that this process is not simply a passive DNA association: tumor DNA is tethered to the RBC surface and can be selectively modulated by glycan remodeling and Toll-like receptor 9 (TLR9) blockade, revealing a regulated mechanism of tumor DNA capture. In lung cancer xenograft models, rtDNA levels correlated with tumor burden, and tumor-specific mutations were detectable in RBCs even when matched plasma samples were negative. Most importantly, in patients with non-small cell lung cancer (NSCLC), RBC-based analysis detected the specific tissue NGS–identified mutation in all patients, and mutant DNA preferentially sequestered with the RBC compartment when compared with plasma. Together, these findings establish RBCs as a biologically regulated reservoir of tumor DNA and reveal a TLR9-dependent mechanism of tumor DNA capture. By expanding the circulating tumor DNA landscape beyond plasma, this work provides a foundation for exploiting the RBC compartment for sensitive cancer detection and disease monitoring.

## Results

### RBC-bound tumor DNA provides a detectable and stable signal

We and others have previously demonstrated that RBCs co-cultured with cancer cell lines can acquire tumor-derived genetic material(*5, 14*). To validate and extend these findings, we first replicated our previous results across three lung cancer cell lines with distinct mutations. Consistent with our prior findings, RBCs incubated with H1975, H358, and A549 cells acquired detectable levels of *EGFR* L858R, *KRAS* G12C, and *KRAS* G12S mutant DNA, respectively. In contrast, these mutations were undetectable in untreated RBCs isolated from healthy donors (Fig. 1A-C). To determine whether tumor-derived DNA acquisition occurs preferentially in reticulocytes as suggested by recent studies(*15*), RBCs depleted of reticulocytes were co-cultured with A549 cells. Reticulocyte-depleted (CD71^-^) RBCs demonstrated similar acquisition of tumor-derived DNA, confirming that mature RBCs are capable of acquiring tumor DNA (Supp. Fig. 1A). We next examined the recoverability of rtDNA following cryopreservation. rtDNA remained stable at -20°C for 7 weeks after co-culture, with minimal degradation observed across RBC samples from five independent healthy donors (Supp. Fig. 1B).

**Figure 1.**
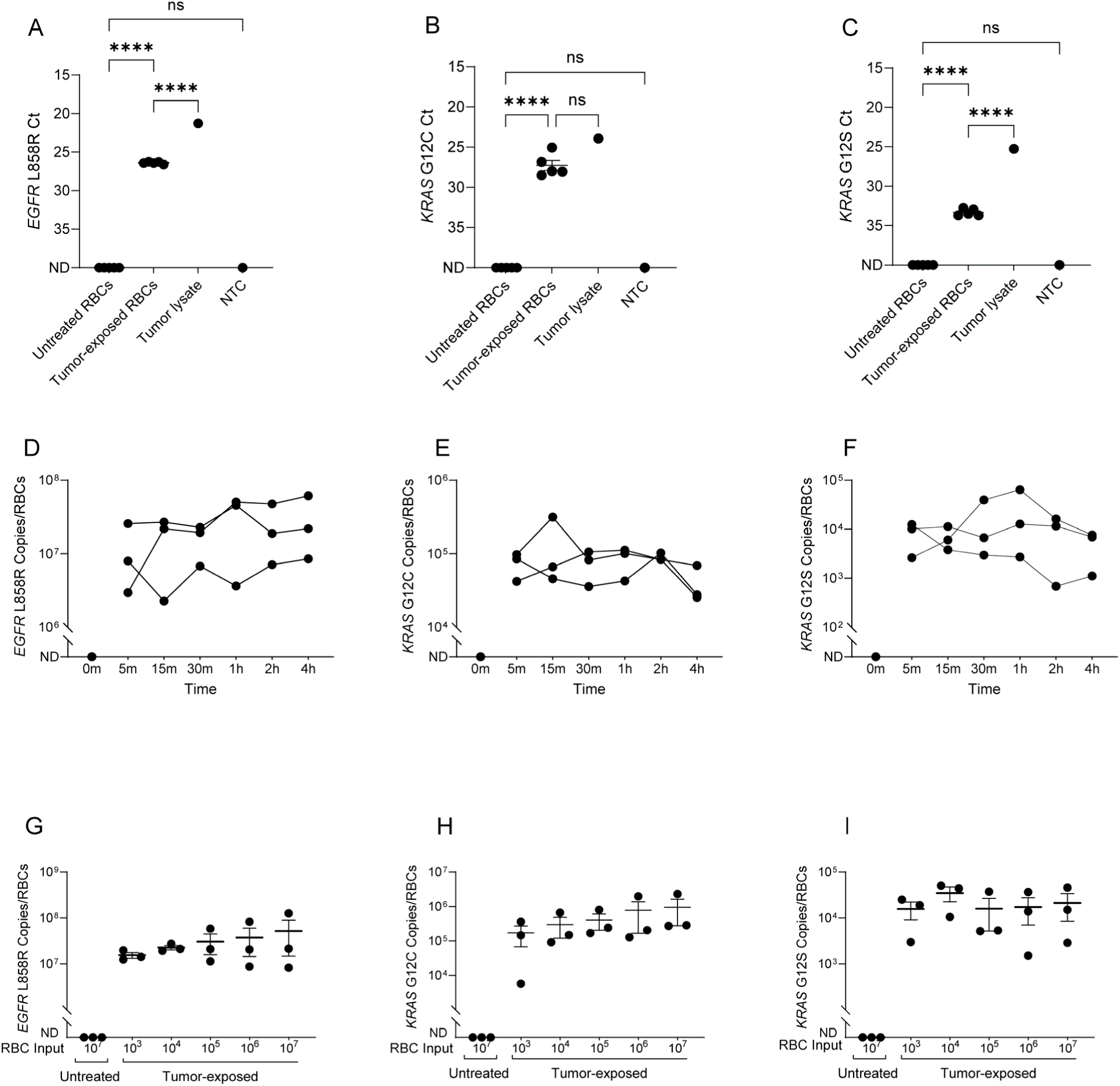
RBCs acquire tumor-derived DNA. **(A–C)** qPCR detection of *EGFR* L858R, *KRAS* G12C, and *KRAS* G12S mutations in RBCs from healthy donors (HDs) following co-culture with H1975, H358, and A549 cells, respectively. Untreated RBCs showed no detectable mutant signal. NTC: no-template control. n=5 HDs, at least 3 independent experiments. Statistical analysis performed with one-way ANOVA with Tukey’s multiple comparisons test. ND: Not Detected. ns: Not significant. **(D-F)** Time-course analysis of tumor DNA acquisition by RBCs following co-culture with H1975, H358, or A549 cells for 5 min to 4 h, with mutant DNA detected at all time points examined. n=3 HDs, 3 independent experiments. No statistical comparisons were performed as signal was undetectable at 0 min and robustly detected at all subsequent time points. **(G-I)** Effect of RBC input on tumor DNA detection, demonstrating detectability at as low as 1,000 RBCs across all three models. n=3 HDs, 3 independent experiments. No statistical comparisons were performed as signal was undetectable with untreated RBCs and robustly detected with tumor-exposed RBCs at all subsequent inputs. Data are presented as mean ± SEM. For all Ct data, reverse axes are presented with a lower Ct denoting greater signal detected. ****P<0.0001.

We next investigated how rapidly RBCs can acquire tumor DNA and the minimum RBC input required for detection. RBCs were co-cultured with each of the three cancer cell lines and harvested at multiple early time points. DNA was extracted followed by quantitative PCR (qPCR). A standard curve was generated using six serial 10-fold dilutions of known concentration of purified cancer lysate amplicon, which were run in parallel with experimental samples on the same qPCR plate to generate copy numbers (Supp. Fig. 1C). Consistent with previous reports(*14*), RBCs rapidly acquired tumor DNA, and rtDNA was detectable as early as 5 min after initiation of co-culture with all cancer cell lines (Fig. 1D-F). In parallel, we systematically varied RBC input to determine the minimum number of cells required for robust tumor DNA detection. Cancer cells were co-cultured with increasing numbers of RBCs, and mutation detection was assessed by qPCR. Remarkably, rtDNA remained detectable at inputs as low as 1,000 RBCs across all three cell lines (Fig. 1G-I). Collectively, these results demonstrate that RBCs can rapidly acquire, sequester, and stably retain tumor-derived DNA.

### Surface localization of tumor DNA on tumor cells and RBCs

We asked whether RBC-mediated tumor DNA acquisition required close proximity to tumor cells. Tumor DNA was not detected on RBCs when co-culture was performed using 0.4 μm transwell inserts. In contrast, tumor DNA acquisition was readily observed under direct co-culture conditions, indicating that acquisition requires close cellular proximity (Supp. Fig. 1D-F). These findings argue against passive uptake of freely diffusible tumor DNA and instead support a localized transfer or capture process at the tumor cell–RBC interface. We next examined whether tumor cells harbor extracellular DNA on their surface. A549 cells were stained with the cell-impermeable nucleic acid dye Sytox under non-permeabilizing conditions. Sytox-positive populations were detected, consistent with the presence of extracellular surface-associated nucleic acids (Fig. 2A). Interestingly, two populations with low (Sytox^Lo^) and high (Sytox^Hi^) green fluorescence were observed, suggesting differences in the abundance or distribution of surface-associated nucleic acids. To determine whether the Sytox signal originated from extracellular DNA, tumor cells were treated with DNase I. DNase treatment markedly reduced the Sytox^Hi^ population, indicating that a significant portion of the detected surface-associated nucleic acids on tumor cells consisted of extracellular DNA (Fig. 2A). These findings demonstrate that tumor cells carry DNase-sensitive extracellular DNA on their surface. We next asked whether RBCs could acquire this surface-bound tumor DNA. A549 cells labeled with Sytox were co-incubated with RBCs for 2 h, after which the remaining tumor cells were analyzed for surface DNA staining. Co-incubation with RBCs resulted in a significant reduction in fluorescence compared with tumor cells cultured alone, and this decrease was most pronounced in the Sytox^Hi^ population of A549 cells (Fig. 2A-C). Data were replicated with another cell line, H358, and showed a similar localization of tumor DNA on the surface of cancer cells that was sensitive to DNase treatment and to RBC co-incubation (Supp. Fig. 1G-I). Together, these findings support a model in which RBCs bind and acquire extracellular DNA present on the surface of tumor cells.

**Figure 2.**
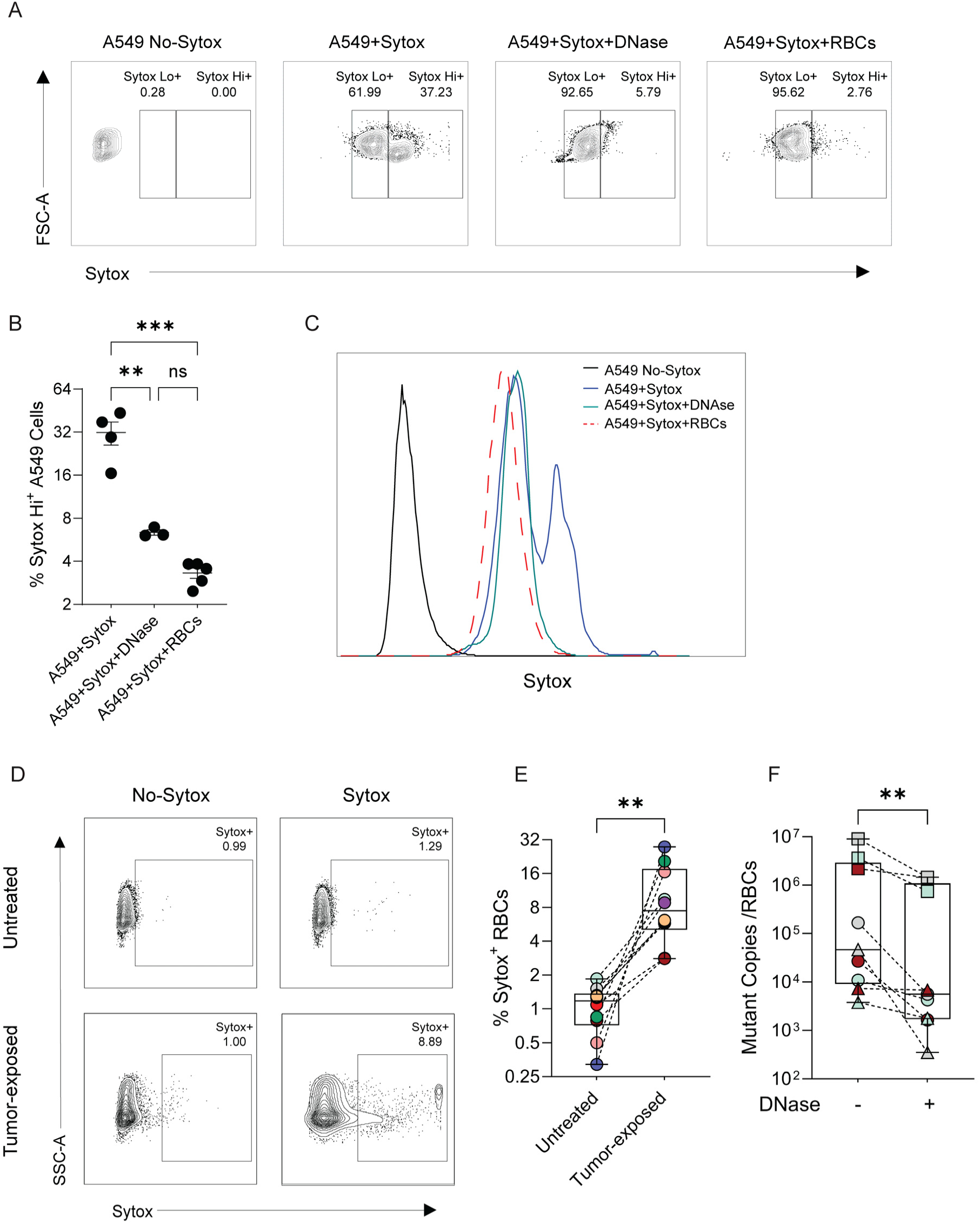
Surface localization of tumor DNA on tumor cells and RBCs. **(A)** Representative contour plots showing Sytox staining of A549 cells under control conditions and following co-incubation with RBCs or DNase treatment. n=5 HDs, one representative experiment shown. Sytox^Hi^: Sytox-high cells; Sytox^Lo^: Sytox-low cells. **(B)** Quantification of Sytox^Hi^ A549 cells following Sytox staining in the presence or absence of RBCs or DNase treatment. n=5 HDs, 3 independent experiments. Data are presented as mean ± SEM. Statistical analysis was performed using one-way ANOVA with Tukey’s multiple comparisons test. **(C)** Representative histograms of Sytox fluorescence corresponding to the conditions shown in (A). **(D)** Representative contour plots showing Sytox staining of untreated RBCs and RBCs following co-culture with tumor cells. n=10 HDs, one representative experiment shown. **(E)** Quantification of Sytox-positive RBCs following co-culture with tumor cells. n=10 HDs, 6 independent experiments. Data are presented as mean ± SEM. Statistical analysis was performed using the paired t-test. **(F)** DNase treatment of RBCs following co-culture with H1975, H358, or A549 cells significantly reduced recoverable tumor DNA. Squares represent *EGFR* L858R, circles represent *KRAS* G12C, and triangles represent *KRAS* G12S. Each color represents an individual RBC donor; n=3 HDs, 3 independent experiments. Data are presented as mean ± SEM. Statistical analysis was performed using the Wilcoxon matched-pairs signed-rank test. **P<0.01, ***P<0.001.

Some reports have suggested that RBC-bound DNA can be localized intracellularly, and that mature RBCs contain long DNA fragments internalized through apoptotic bodies(*14, 16*). To assess if RBC-bound DNA can be present on the RBC cell surface, flow cytometry was performed using Sytox. Tumor-exposed RBCs exhibited a significant increase in fluorescence compared with untreated controls, confirming the presence of cell-surface-bound nucleic acids (Fig. 2D-E), although only a relatively small population was positive. To further validate this observation, we tested whether rtDNA was sensitive to enzymatic degradation. Intact, non-permeabilized RBCs were co-cultured with tumor cells from 3 cell lines and subsequently treated with DNase. DNase treatment attenuated the recoverable tumor DNA from RBCs (*EGFR* L858R, *KRAS* G12C, and *KRAS* G12S mutants detected by qPCR), confirming surface localization of tumor DNA (Fig. 2F), similar to mitochondrial DNA (Supp. Fig. 1J). Collectively, these data indicate that RBC-mediated tumor DNA acquisition occurs through local capture of surface-associated extracellular DNA at the tumor cell–RBC interface rather than by passive uptake of freely diffusible DNA.

### Tumor DNA acquisition by RBCs is tunable and dependent on TLR9

Interestingly, across all healthy donors, only a small subset of RBCs demonstrated Sytox positivity following incubation with tumor cells, revealing heterogeneity in DNA acquisition both between donors and within individual donor RBC populations (Fig. 2E). We have previously demonstrated that RBCs express the DNA-sensing receptor TLR9 on their surface and can bind host and pathogen DNA(*1–3*). The donor-dependent variability in tumor DNA acquisition observed here mirrored our prior observations of RBC-TLR9-mediated CpG binding, leading us to hypothesize that tumor DNA acquisition may likewise occur through a TLR9-dependent mechanism. To test this, RBCs were treated with either isotype control IgG or an anti-TLR9 antibody (which we have previously shown interferes with DNA-induced RBC senescence(*3*)) prior to co-culture with tumor cells. We observed a significant decrease in *KRAS* G12S mutant copies acquired by RBCs following TLR9 blockade, indicating that tumor DNA acquisition is mediated, in part, by TLR9 (Fig. 3A), supporting our data that the DNA is bound on the RBC surface.

**Figure 3.**
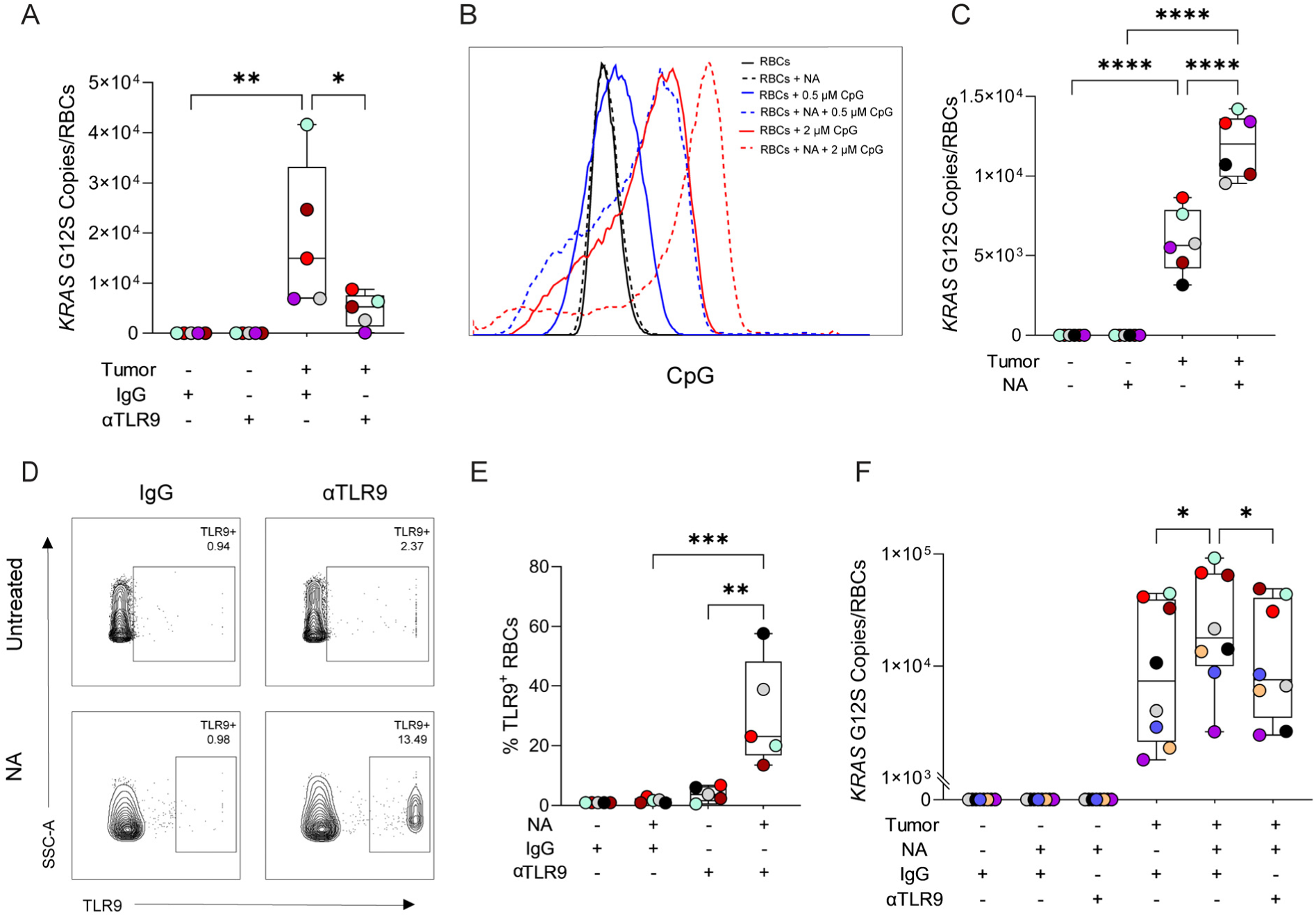
Tumor DNA acquisition by RBCs is tunable and dependent on a TLR9-associated mechanism. **(A)** TLR9 blockade attenuated RBC acquisition of tumor-derived DNA. n=5 HDs, 3 independent experiments. Statistical analysis was performed using one-way ANOVA with Šidák’s multiple comparisons test. **(B)** Neuraminidase treatment enhanced RBC acquisition of exogenous DNA. Two independent experiments are shown, each performed using 0.5 μM and 2 μM CpG. NA: Neuraminidase. **(C)** NA treatment increased *KRAS* G12S tumor DNA acquisition following co-culture with A549 cells. n=6 HDs, 3 independent experiments. Statistical analysis was performed with one-way ANOVA with Šidák’s multiple comparisons test. **(D)** Representative contour plots demonstrating surface TLR9 expression on RBCs following NA treatment. **(E)** Quantification of surface TLR9 expression following NA treatment. n=5 HDs, 5 independent experiments. Statistical analysis was performed using one-way ANOVA followed by Šidák’s multiple comparisons test. **(F)** NA-treated RBCs exhibited enhanced *KRAS* G12S tumor DNA acquisition following co-culture with A549 cells, which was attenuated by anti-TLR9 treatment. n=8 HDs, 4 independent experiments. Statistical analysis was performed using one-way ANOVA followed by Šidák’s multiple comparisons test. Data are presented as mean ± SEM unless otherwise indicated. *P<0.05, **P<0.01, ***P<0.001, ****P<0.0001.

Given that the RBC surface is densely coated with sialylated glycans that regulate receptor accessibility and cell–environment interactions, we investigated whether remodeling of the RBC glycocalyx could modulate tumor DNA acquisition. Neuraminidase (NA) treatment removes terminal sialic acid residues from surface glycoconjugates, we thus examined the effects of NA treatment on RBC-mediated DNA acquisition. Desialylated RBCs showed an increased capacity to acquire DNA when incubated with synthetic CpG oligonucleotides (Fig. 3B), or when co-cultured with A549 cancer cells (Fig. 3C). Because surface remodeling altered RBC-mediated DNA acquisition, we hypothesized that RBC-DNA receptor availability may be increased following desialylation; thus, we subsequently evaluated the impact of NA treatment on RBC surface TLR9 detection. Flow cytometric analysis demonstrated increased surface TLR9 detection following desialylation (Fig. 3D-E). We then investigated whether TLR9 blockade would attenuate NA-enhanced tumor DNA uptake by RBCs. Consistent with our previous observations, NA-treated tumor-exposed RBCs exhibited significantly increased rtDNA compared with untreated tumor-exposed RBCs. Importantly, this augmentation of DNA acquisition was attenuated when desialylation was combined with TLR9 blockade, further supporting a TLR9 and RBC surface-dependent mechanism of DNA acquisition (Fig. 3F). Collectively, these findings demonstrate that tumor DNA is localized on the surface of RBCs and that RBC-mediated DNA capture can be enhanced by surface remodeling. Moreover, these data suggest that tumor DNA is acquired through a receptor-mediated process involving TLR9.

### RBCs acquire tumor DNA in vivo

We next evaluated whether RBC-mediated tumor DNA acquisition was observable in a murine system. Similar to human RBCs, murine RBCs acquired tumor DNA following co-culture with human tumor cells (Supp. Fig. 2A). To evaluate the feasibility of RBC-mediated tumor DNA for liquid biopsy, RBCs from tumor-bearing mice were assayed for mutant DNA. Immunodeficient NOD Scid Gamma (NSG) mice were injected via tail vein with A549-NY-ESO-luc human lung adenocarcinoma cells which contain a *KRAS* G12S mutation. Tumor establishment and progression were monitored using an *in vivo* imaging system (IVIS), which confirmed successful engraftment of tumor cells (Fig. 4A-B). Peripheral blood samples were collected up to 3 weeks post-injection to assess the ability of RBCs to acquire tumor DNA. RBCs were isolated, DNA was extracted, and qPCR was performed to detect mutant *KRAS* G12S DNA. *KRAS* G12S copies were readily detected on RBCs of A549-injected mice, with a significant correlation observed between RBC-bound mutant DNA and tumor burden as measured by bioluminescence (Fig. 4C).

**Figure 4.**
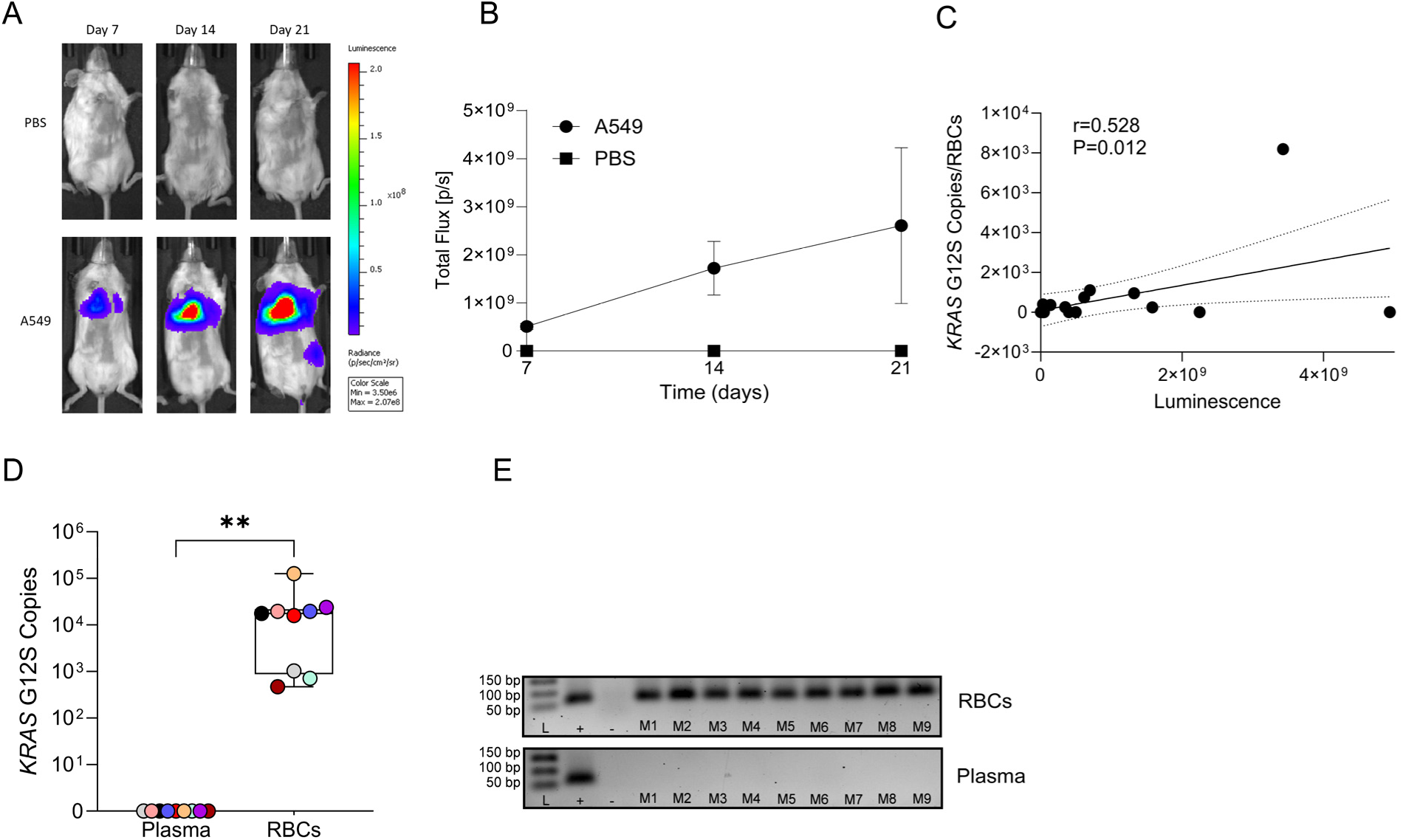
RBCs acquire tumor DNA with greater detection rates than plasma in murine lung cancer models. **(A)** Representative IVIS imaging confirming tumor establishment in NSG mice following intravenous injection of A549-NY-ESO-luc cells. **(B)** Quantification of total flux in tumor cell-injected and PBS-treated mice (n=10). A549 cells produced significantly higher bioluminescent signal compared with PBS controls on day 7 (P=0.0043), with a trend toward significance on day 14 (P=0.11), that did not reach significance on day 21 due to increased variance in tumor burden (P=0.46), two-way RM ANOVA with Šidák correction. **(C)** Correlation analysis between tumor burden and RBC-associated *KRAS* G12S mutant copy number (n=10). **(D)** Comparative qPCR analysis of matched RBC and plasma samples collected from NSG mice 2 months after subcutaneous injection of A549 cells (n=9). Statistical analyses were performed using the Wilcoxon test. **(E)** Representative gel electrophoresis confirming *KRAS* G12S amplicons in RBC-derived DNA, with absent signal in matched plasma samples. The gel image was processed using ImageJ. L: Ladder; M: Mouse; (+): A549 cell lysate; (−): no-template control. **P<0.01.

After confirming that RBCs can acquire tumor DNA in an A549-mouse model of lung cancer, we next investigated whether RBCs could serve as a superior substrate for tumor DNA detection compared with plasma. To address this, a subcutaneous xenograft model was established in NSG mice using A549 cells(*17*), and tumors were allowed to develop for two months prior to sample collection. Peripheral blood was collected, and matched plasma and RBC fractions were isolated for comparative analysis. DNA was extracted from each fraction (RBC vs plasma), followed by qPCR targeting mutant *KRAS* G12S. *KRAS* G12S mutant DNA was detectable on RBCs from all animals, in contrast to a complete absence in plasma (Fig. 4D-E).

### Tumor-derived DNA is detectable on RBCs in patients with NSCLC and is concordant with matched tumor tissue

We next asked whether these findings could be translated to a clinical setting. We recruited 9 patients with late-stage (LS) NSCLC (Table 1) and 3 patients with early-stage (ES) NSCLC (Table 2), whose tumor tissue genotyping by NGS identified *EGFR* L858R mutations in LS1–4, *KRAS* G12R mutations in LS5–6, and *KRAS* G12C mutations in LS7–9 and ES1–3 patients. Peripheral blood samples were collected from all patients, plasma was collected and RBCs were isolated prior to DNA extraction and molecular analysis by qPCR and droplet digital PCR (ddPCR). qPCR analysis demonstrated consistent and enhanced detection of tumor-derived DNA in the RBC fraction across all patients, whereas no mutant signal was detected in RBCs from healthy donors (qPCR analysis of RBCs from patients ES1–3 has also been previously reported(*5*)) (Fig. 5A-B). We next performed ddPCR analysis of patient-derived RBC DNA to further characterize the tumor-associated DNA signal. A detectable mutant signal was observed in RBCs from all patients, together with the corresponding wild-type (WT) signal, demonstrating that RBCs acquired both mutant and WT DNA irrespective of the underlying tumor mutation (Fig. 5C). To directly compare the capacity of RBCs and plasma to capture tumor DNA, equal volumes of RBCs and plasma (5 μL each) were processed for DNA extraction and ddPCR. Across both WT and mutant targets, tumor DNA was detected at significantly higher levels in RBCs than in plasma (Fig. 5D-E). Across the cohort, ∼30% of the DNA detected in paired RBC and plasma samples carried the tumor-specific mutant allele, whereas ∼70% carried the corresponding WT allele; patient-level analysis revealed that >99% of this detectable tumor-specific mutant DNA was RBC-bound rather than plasma-derived (Fig. 5F-G). Notably, rtDNA, detected by both qPCR and ddPCR, recapitulated the tumor-specific mutations identified by tissue NGS in all 12 patients. By comparison, among the six patients with matched tissue NGS and results from a commercial plasma-based liquid biopsy assay, the corresponding driver mutation was detected in only three patients (50%) (Fig. 5H and Table 1). Collectively, these findings extend our preclinical observations to patient samples and demonstrate that tumor-derived DNA associated with RBCs can be detected at levels sufficient to identify tumor-specific mutations, including mutations that were not detected with commercial plasma-based liquid biopsy platforms.

**Figure 5.**
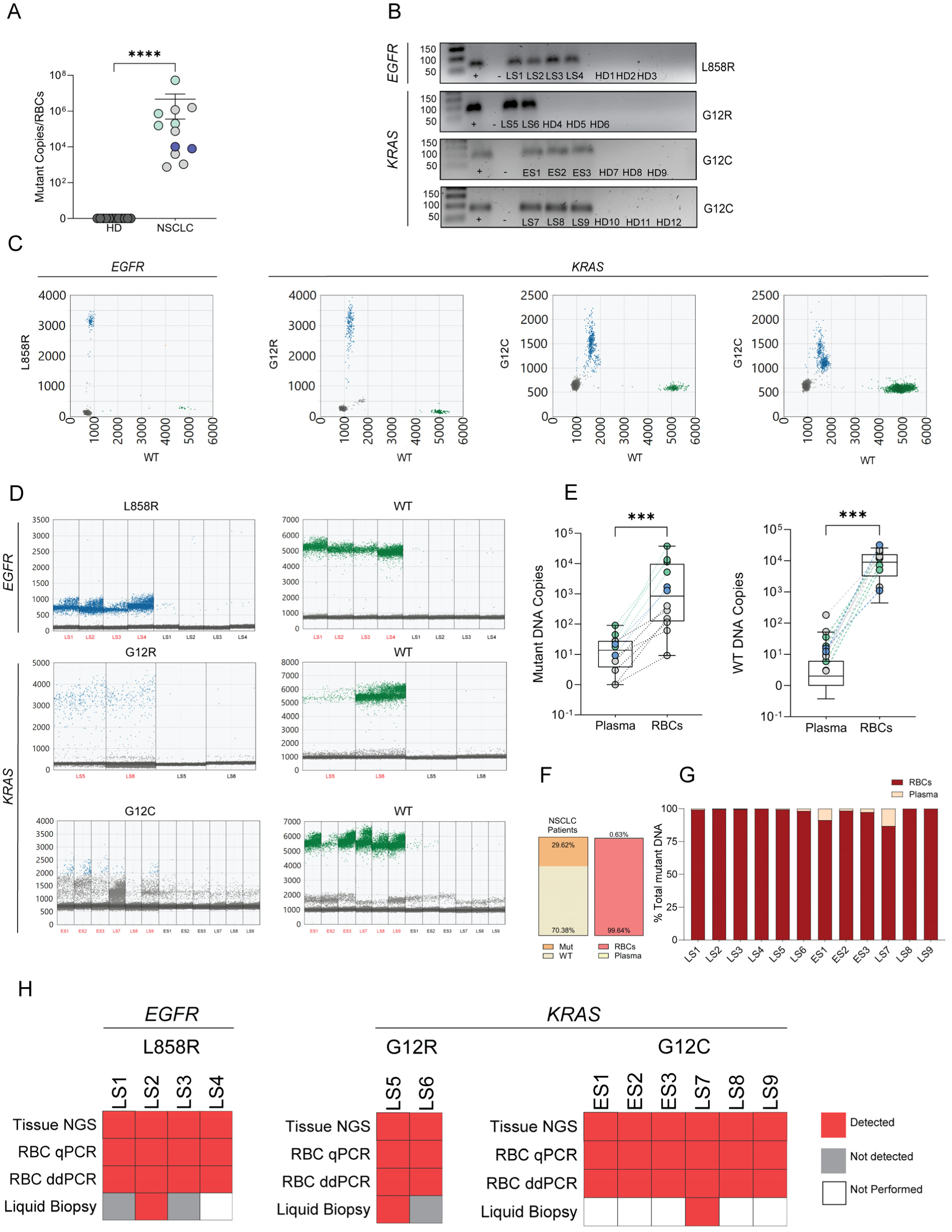
RBCs detect tumor DNA in patients with lung cancer with concordance to tissue NGS. **(A)** qPCR detection of tumor-associated mutations in RBCs from patients with NSCLC, including *EGFR* L858R (green), *KRAS* G12R (blue), and *KRAS* G12C (grey); n=12 NSCLC patients and 12 HDs. Data are represented as mean ± SEM, statistical analysis performed using the Mann Whitney test. **(B)** Representative gel electrophoresis comparing amplicon detection in healthy donor versus NSCLC patient RBC-derived DNA, demonstrating tumor-specific bands in patient samples and absence of signal in controls. n=12 NSCLC patients and 12 HDs. The gel image was processed using ImageJ. Gels represent *EGFR* L858R, *KRAS* G12R, *KRAS* G12C ES, and *KRAS* G12C LS patients respectively from top to bottom. LS: Late-Stage NSCLC patient; ES: Early-Stage NSCLC patient; HD: Healthy Donor; (+) tumor cell lysate; (−) no-template control. **(C)** 2D droplet amplitude plots showing individual ddPCR droplets based on fluorescence intensity for the WT and mutant signals. The x-axis represents the WT signal, while the y-axis represents the mutant signal (*EGFR* L858R, *KRAS* G12R, and *KRAS* G12C). Blue droplets represent mutant-positive droplets, whereas green droplets represent WT-positive droplets. One representative LS patient was graphed per mutation (LS2 for *EGFR* L858R, LS5 for *KRAS* G12R, and LS9 for *KRAS* G12C), and one representative ES patient for G12C (ES2). **(D)** 1D droplet amplitude plots showing the detection of the tumor mutation in plasma and RBC-DNA samples. Each droplet is represented based on its fluorescence amplitude, with mutant-positive and WT droplets separated according to the assay-defined fluorescence thresholds. Matching RBC and plasma samples from each patient are labeled in red and black respectively with the corresponding patient number. n=12 NSCLC patients. LS: Late-Stage NSCLC patient; ES: Early-Stage NSCLC patient. **(E)** Quantification of mutant- and WT-positive droplets in plasma and RBC-DNA from patients with NSCLC. Statistical analyses were performed using the Wilcoxon test. *EGFR L858R*, *KRAS* G12R, and *KRAS* G12C patients are represented by green, blue, and grey circles respectively. **(F)** Left pie chart shows mutant versus wild-type DNA detected across all 12 patients in combined RBC and plasma samples, with mutant DNA comprising ∼30% of the total detected DNA. The right pie chart shows % detectability of mutant DNA on RBCs versus plasma. **(G)** Percentage of total mutant DNA detected in RBCs versus plasma for each patient. **(H)** Heat map showing detection of tumor-associated mutations in RBCs from NSCLC patients, including *EGFR* L858R, *KRAS* G12R, and *KRAS* G12C, n=12. Comparison of mutation detection by tissue NGS, RBC qPCR, RBC ddPCR, and commercially available liquid biopsy. LS: Late-Stage NSCLC patient; ES: Early-Stage NSCLC patient. ***P<0.001, ****P<0.0001.

**Table 1.** Late-Stage Patient Demographics and Liquid Biopsy Results.

| Patient ID | Age at Dx | Sex | Race | Stage | Solid Tumor Biopsy Site | Solid Tumor NGS * | Solid Tumor VAF | rtDNA PCR | rtDNA Concor -dance | Liquid Biopsy Concor -dance | Liquid Biopsy NGS VAF<sup>#</sup> | Liquid Biopsy Tumor Fraction |
| --- | --- | --- | --- | --- | --- | --- | --- | --- | --- | --- | --- | --- |
| LS1 | 75 | F | White | IVA | Lung | *EGFR* L858R | 45% | *EGFR* L858R | Yes | No | Not Reported | Not Reported |
| LS2 | 68 | F | White | IVB | Pleural Fluid | *EGFR* L858R | 12% | *EGFR* L858R | Yes | Yes | 0.3% | 0.02% |
| LS3 | 83 | F | White | IVA | Lung | *EGFR* L858R | 37% | *EGFR* L858R | Yes | No | N/A | 0.06% |
| LS4 | 66 | M | White | IVA | Lung | *EGFR* L858R | 49% | *EGFR* L858R | Yes | Not Performed | N/A | N/A |
| LS5 | 59 | M | White | IVB | Pleural Fluid | *KRAS* G12R | 6% | *KRAS* G12R | Yes | Yes | 2.2% | 0.4% |
| LS6 | 81 | M | White | IVA | Lung | *KRAS* G12R | 38% | *KRAS* G12R | Yes | No | N/A | 0.06% |
| LS7 | 78 | M | Black | IVA | Pleura | *KRAS* G12C | 40% | *KRAS* G12C | Yes | Yes | 0.07% | 0.10% |
| LS8 | 78 | M | White | IVA | Lung | *KRAS* G12C | 49% | *KRAS* G12C | Yes | Not Performed | N/A | N/A |
| LS9 | 75 | M | Black | IVA | Pleura | *KRAS* G12C | 14% | *KRAS* G12C | Yes | Not Performed | N/A | N/A |

Age at Dx: Age at diagnosis
F: Female/ M: Male
\*Solid tumor NGS completed using an in-house comprehensive NGS platform
<sup>#</sup>Liquid biopsy NGS completed using the Guardant 360 NGS platform
N/A: Not Applicable

**Table 2.** Early-Stage Patient Demographics.

| **Patient ID** | **Age at Dx** | **Sex** | **Stage** | **Histology** | **Solid Tumor NGS \*** |
| --- | --- | --- | --- | --- | --- |
| ES1 | 77 | F | IA | Adenocarcinoma | *KRAS* G12C |
| ES2 | 70 | F | IIB | Adenocarcinoma | *KRAS* G12C |
| ES3 | 66 | F | IA | Adenocarcinoma | *KRAS* G12C |

Age at Dx: Age at diagnosis
F: Female
\*Solid tumor NGS completed using an in-house comprehensive NGS platform

## Discussion

In this study, we identify molecular determinants of tumor DNA binding to RBCs and establish the relevance of the RBC surface as a distinct compartment for liquid biopsy. Building upon our prior findings that RBCs bind host, pathogen, and tumor-derived nucleic acids(*1–5*), our observations link RBC membrane biology to the retention and detection of tumor DNA. Collectively, these findings extend the current understanding of RBC-associated DNA biology and provide a framework for its potential clinical application in cancer diagnostics.

Our prior work established that RBCs participate in nucleic acid regulation by binding host and pathogen DNA through surface expression of TLR9, resulting in accelerated RBC senescence and altered host responses during sepsis(*2, 3*). In the current study, we extend this paradigm to cancer biology and show that RBCs could acquire and stably harbor tumor-derived DNA across multiple lung cancer cell lines carrying clinically relevant oncogenic mutations, in mouse xenograft models, and in patients with lung cancer. Our findings establish RBCs as a distinct reservoir of tumor-derived DNA with direct relevance to liquid biopsy, offering several important advantages including their exceptional abundance, long circulating lifespan, and lack of a nucleus, which may provide a stable and relatively low-background compartment for extracellular tumor DNA capture. Importantly, our experiments demonstrate complete absence of detectable rtDNA in untreated healthy-donor RBCs, which minimizes concerns regarding contamination or nonspecific amplification that represent persistent challenges in ultra-sensitive liquid biopsy assays(*18*).

Additionally, we observed that rtDNA remained readily detectable and stable even after prolonged storage. We speculate that the RBC membrane may provide a protective microenvironment that stabilizes bound DNA fragments and preserves mutational signal integrity over extended periods. This finding has practical implications for downstream translational workflows, suggesting that archived specimens may remain suitable for retrospective or delayed molecular analyses. This stability represents a potential advantage over plasma ctDNA whose sensitivity remains limited by the scarcity and rapid clearance of tumor-derived DNA from the circulation; the ability of priming agents to transiently attenuate cell-free DNA clearance and thereby increase ctDNA recovery and improve detection of small tumors(*19*) provides direct evidence that analyte availability is a major constraint on the sensitivity of liquid biopsy. These findings underscore the need not only for improved analytical methods, but also for alternative strategies to capture and preserve tumor-derived DNA within the circulation. In this context, our identification of RBCs as a circulating reservoir of tumor-derived DNA provides an alternative approach that leverages a naturally abundant cellular compartment. While the *ex vivo* stability demonstrated here supports the feasibility of clinical translation including the use of archived or biobanked specimens for rtDNA detection, the *in vivo* kinetics of rtDNA remain largely undefined, necessitating dedicated studies to determine its half-life and clearance dynamics.

RBCs are increasingly recognized as active immunomodulatory cells that may participate in immune surveillance(*20, 21*). Because tumors are amongst the most vascularized tissues due to angiogenesis, and RBCs are the most abundant and long-lived cell in circulation, there are innumerable opportunities for RBC contact with tumor cells allowing for transfer of proteins and nucleic acids from tumor to RBC(*22–25*). In this study, we provide evidence that RBCs acquire tumor-derived DNA primarily through direct interaction with tumor cells rather than by passive uptake of freely circulating DNA. Our findings support a model in which RBCs function as scavengers of extracellular tumor DNA at sites of direct tumor cell contact. A key observation from this study is the presence of DNase-sensitive extracellular DNA on the surface of tumor cells. Although cell-surface-associated nucleic acids have received relatively little attention compared with circulating cell-free DNA, some suggest that this phenomenon may represent a conserved feature of malignant cells contributing to the tumor microenvironment by facilitating cell adhesion, migration, signaling, and interactions with surrounding immune cells(*26, 27*). Our data extend this concept by identifying RBCs as a previously unrecognized cell type capable of interacting with this extracellular DNA pool. Rather than functioning solely as oxygen carriers, RBCs may participate in regulating extracellular nucleic acid availability by binding DNA present on the tumor cell surface. By capturing tumor-derived DNA, RBCs may regulate the availability of DNA to immune cells and thereby influence immune responses. As circulating reservoirs of tumor-associated DNA, RBCs could play a role in either promoting immune tolerance or supporting immune surveillance, depending on the biological context.

The mechanism through which tumor DNA becomes associated with the tumor cell surface remains incompletely understood. Tumor cells continuously release DNA through apoptosis, necrosis, and extracellular vesicle secretion(*28*). Following its release, extracellular DNA could remain localized at the cell surface or associated with extracellular vesicles through interactions with histones, DNA-binding proteins, and extracellular matrix components such as proteoglycans, forming an extracellular DNA layer. Under physiological conditions, circulating RBCs are largely confined to the vascular lumen and would be expected to have limited opportunity to directly contact tumor cells. However, many solid tumors develop abnormal vascular architecture characterized by increased permeability, disrupted endothelial barriers, and direct exposure of RBCs to tumor tissue(*29*). In addition, increasing evidence supports the phenomenon of vasculogenic mimicry, whereby highly aggressive tumor cells acquire endothelial-like properties and form vessel-like networks capable of facilitating blood perfusion independently of conventional endothelial-lined vasculature(*30–32*). Within these structures, RBCs may encounter tumor cell surfaces directly, providing a setting for the capture of cell-surface tumor DNA. Our observation that DNA acquisition requires direct cellular proximity is consistent with this model and raises the possibility that vascular mimicry and other forms of abnormal tumor angiogenesis facilitate RBC acquisition of tumor-derived DNA, though further studies will be needed to definitively establish the mechanistic basis for RBC-mediated tumor DNA acquisition.

The extracellular localization of mitochondrial DNA on the mammalian cell surface has been recently reported(*33*), while other reports suggest internalization of tumor DNA by RBCs(*14, 16*). Here, we observe susceptibility of rtDNA to enzymatic digestion by DNase, indicating that a substantial proportion of rtDNA is externally bound to the RBC surface rather than internalized. Notably, rtDNA is detected in only a subset of RBCs, suggesting that tumor DNA acquisition is not a uniform property of all erythrocytes. Emerging evidence indicates that RBCs are functionally heterogeneous, with distinct subpopulations differing in surface receptors, bound nucleic acids, and immune-related functions(*3, 21*). We have previously demonstrated heterogeneous TLR9 expression on RBCs and showed that RBCs participate in host responses to circulating DNA(*1–3*). RBC-TLR9 has been implicated in the regulation of mitochondrial and microbial DNA, particularly during inflammatory states, such as sepsis and critical illness, and has more recently been implicated in the pathogenesis of multiple sclerosis(*1, 3, 34*). Upon binding DNA, RBCs undergo membrane remodeling and are subsequently cleared by macrophages, suggesting that they function as sacrificial carriers that facilitate the removal of inflammatory nucleic acids from the circulation. Given the known interindividual variability in RBC-TLR9 expression, we hypothesized that TLR9 contributes to tumor DNA acquisition. Indeed, TLR9 blockade attenuated tumor DNA acquisition, implicating TLR9 as a mediator of rtDNA capture.

RBC membrane glycoproteins are heavily coated with sialic acids that contribute to surface charge, immune evasion, and receptor masking(*35–39*). Desialylating RBCs markedly enhanced tumor DNA acquisition. Variation in RBC membrane glycoproteins and sialic acid composition occurs during RBC aging and across blood groups, and is known to modulate interactions with heterotypic cells (including B cells and neutrophils) and alter RBC-pathogen interactions(*40–44*). During inflammatory states, such as bacterial infection and sepsis, RBC desialylation has been reported and recent data suggest these alterations may contribute to heterotypic interactions with immune cells(*41, 43, 45*). Thus, variable RBC sialylation due to differences in blood group antigen density on the RBC surface or inflammatory state may modulate receptor accessibility and contribute to donor-specific differences in rtDNA acquisition capacity(*44*). Under basal conditions, RBC-TLR9 accessibility remains limited; however, consistent with our previous observations(*3*), RBC membrane alterations, now including desialylation, enhance surface detection and RBC-mediated DNA capture. We speculate that sialylation of the RBC surface not only serves to provide the negative charge and known repulsive effect of RBCs but also serves to limit RBC-TLR9-mediated immune activation. This is consistent with previous studies demonstrating that loss of RBC sialylation permits B cell and neutrophil activation(*41, 42*). While our data do not exclude participation of additional receptors or DNA internalization, they strongly support a model in which RBC-TLR9 contributes to surface capture and retention of tumor-derived DNA fragments. Our findings also suggest a potential therapeutic opportunity through modulation of the RBC glycocalyx. Sialidases have been investigated as a means to remodel the glycocalyx of cancer cells, increasing the exposure of tumor-associated surface epitopes and enhancing immune recognition(*46*). A similar strategy could potentially be applied to circulating RBCs. Controlled desialylation of RBCs may expose additional membrane components capable of interacting with tumor DNA including TLR9, thereby enhancing the ability of RBCs to function as tumor reservoirs and improving the sensitivity of RBC-based liquid biopsy approaches.

The rapid and durable accumulation of tumor-derived DNA by RBCs suggests that rtDNA represents a novel tool for liquid biopsy. Supporting this premise, tumor-bearing mice demonstrated mutant *KRAS* DNA on RBCs from all mice, while no mutant DNA was observed in matched plasma samples. Consistent with these findings, RBCs from patients with early- and late-stage NSCLC demonstrated robust detection of *EGFR* and *KRAS* mutations identified by tissue mutational analysis, even when they were absent in conventional plasma-based assays. Utilizing rtDNA as an analyte holds the potential to improve the detection of actionable genetic alterations in advanced NSCLC, allowing more eligible patients to benefit from efficacious molecularly directed therapies. A major advantage of rtDNA is its compatibility with rapid and widely accessible molecular assays. In this study, tumor-associated DNA bound to RBCs was readily detected using qPCR and ddPCR, demonstrating that mutant DNA detection can be achieved without relying on NGS. For applications targeting known mutations or tumor-specific markers, these PCR-based approaches offer substantially shorter turnaround times, lower costs, and greater accessibility than sequencing-based methods. Because PCRs are already routinely implemented in clinical and research laboratories, analysis of rtDNA has the potential to provide a scalable and practical liquid biopsy platform that can be readily integrated into existing diagnostic workflows. Although previous studies have provided evidence that RBCs can acquire extracellular DNA, detection of tumor-specific mutations in RBC-derived DNA has remained limited; ddPCR detection of the *EGFR* L858R mutation in clinical rtDNA was not achieved in the tested mutant sample, despite readily detectable mutant copies in matched plasma cell-free DNA(*16*). Similarly, low-abundance *BRAF* V600E signal was detected in RBCs following exposure to tumor-conditioned media, in marked contrast to the robust signal observed in platelets(*47*). These limited detection rates may reflect RBC-based technical challenges, including incomplete RBC purification, limited DNA recovery, differences in initial blood volumes, and heme-associated inhibition of downstream amplification. Here, we optimized RBC isolation and downstream applications to improve recovery and minimize technical interference, including RBC purification, optimization of sample input, DNA concentration and dilution to mitigate heme-associated inhibition of downstream ddPCR (Supp. Fig. 2B). These methodological improvements enabled reproducible detection of tumor-specific mutant alleles in highly purified RBC-derived DNA, supporting the use of RBCs as a source of tumor-associated DNA that can be interrogated by mutation-specific assays.

Several limitations should be acknowledged. First, our patient cohort was small and consisted primarily of advanced-stage lung cancer patients. Larger prospective studies will be required to determine diagnostic sensitivity, specificity, and clinical utility of RBCs in the early detection of lung cancer. Second, the molecular analyses performed here were limited to mutation detection on RBCs contained within less than 5 μL of blood; the diagnostic potential may be substantially greater when larger RBC volumes are analyzed. Additionally, we intentionally limited our analysis to PCR-based methods to test the feasibility of a relatively cost-effective, simple, and rapid method of RBC-based liquid biopsy. Given the increasing importance of NGS, DNA methylation, and fragmentomic approaches to interrogating ctDNA in precision oncology, future studies will explore whether RBC-derived tumor DNA can support these applications. Large-scale clinical validation will be required to determine whether RBC-based liquid biopsy approaches can complement or ultimately supplant current plasma ctDNA technologies.

Our data demonstrate that RBCs can serve as highly sensitive reservoirs of tumor DNA, highlighting their potential role in liquid biopsy applications. RBCs rapidly acquire tumor DNA through partially TLR9-dependent mechanisms, retain stable mutational signals over time, and detect oncogenic mutations in both murine models and lung cancer patients even when plasma-based assays failed to detect them. These findings establish rtDNA as a novel platform for the identification of actionable genomic abnormalities and disease monitoring, fundamentally redefining the landscape of liquid biopsy and raising the possibility of applications in cancer screening and early detection. Thus, tumor DNA capture and retention establish RBCs as tumor sentinels.

## The authors declare no acknowledgments

### Funding

This work was supported by funding from the Sponsored Research Agreement (SRA) between the University of Pennsylvania and Senticell (NSM), Abramson Cancer Center Support Grant (NSM) from the NCI, P30 CA016520, and NCI R41 CA314866 (Senticell, NSM, JP).

### Author contributions

Experiments were conceived and designed by E.E. and N.S.M. Experiments were performed by E.E., H.R., P.A., E.N.O., and A.K. Data were analyzed by E.E., H.R., J.P., P.A., E.N.O., A.K., J.C.T., S.M.A., and N.S.M. B.A.B., M.G.S.S., and C.L. provided biosamples and phenotypes from cancer patients. S.M.A. and J.P. provided vital reagents. E.E. and N.S.M. wrote the paper.

### Competing interests

N.S.M. is an inventor on the following patent applications related to this work: “Methods for Detection of Pathogenic Infections Using Red Blood Cell-Containing Patient Samples” (US, PCT, and national phase applications); “Detection of Tumor-Derived Nucleic Acids Using Red Blood Cells” (N.S.M. and J.C.T.; US, PCT, and national phase applications); and “Nucleic Acid Delivery Using Direct Binding of Nucleic Acids to RBCs” (US, PCT, and national phase applications). A provisional patent application relating to aspects of the technology described in this manuscript has been filed. N.S.M. is a founder, equity holder, and scientific advisor of Senticell and reports research funding from Senticell. S.M.A. is a scientific advisor for Senticell. B.A.B. has received consulting fees from Johnson & Johnson and honoraria from MJH Life Sciences. M.G.S.S. and J.P. report research funding from Senticell.

### Data, code, and materials availability

All data are available in the main text or the supplementary materials. All materials used in this study are commercially available from the sources indicated in the Materials and Methods.

## Materials and Methods

### RBC processing and isolation

#### Isolation of RBCs from human whole blood

Whole blood samples obtained from healthy donors or patients with cancer were centrifuged at 3,000×g for 10 min. Following centrifugation, plasma was carefully collected without disturbing the buffy coat or packed RBCs (pRBCs) layer. Buffy coat was carefully aspirated and discarded to minimize contamination of the RBC fraction.

For magnetic purification of RBCs, 30 µL of pRBCs was resuspended in 70 µL MACS buffer (Miltenyi Biotec). Unless otherwise specified, reagents and samples were maintained at 4°C throughout the isolation procedure. Anti-CD235a (glycophorin A) magnetic beads (15 µL; Miltenyi Biotec) were added to each sample, mixed thoroughly by pipetting, and incubated for 15 min at 4°C on a nutator. Following incubation, cells were washed with 1 mL MACS buffer and centrifuged at 300×g for 5 min. The supernatant was removed, and the cell pellet was resuspended in 500 µL MACS buffer.

Magnetic separation was performed using LS Columns (Miltenyi Biotec) positioned in a magnetic separator. Columns were equilibrated with 3 mL MACS buffer before sample application. The labeled cell suspension was loaded onto the column and allowed to pass through completely, followed by two washes with 3 mL MACS buffer. The column was then removed from the magnetic field and placed over a 15-mL conical tube. Bound cells were eluted by applying 5 mL MACS buffer using the supplied plunger. Eluted cells were centrifuged at 800×g for 5 min, washed once with 5 mL MACS buffer, and centrifuged again at 800×g for 5 min. The final RBC pellet was resuspended in 1 mL MACS buffer and counted. Cell suspensions were subsequently adjusted to 10⁸ RBCs/mL.

For experiments requiring DNA extraction or storage, 100-µL aliquots containing 10⁷ RBCs were transferred to DNA LoBind tubes (Fisher Scientific), centrifuged, and resuspended in 100 µL DNA/RNA Shield. Depending on the experimental application, RBCs were alternatively resuspended in PBS (Corning) or appropriate culture medium for subsequent incubation experiments. Samples designated for storage were maintained at −80°C until further processing.

#### Isolation of pRBCs for direct plasma-to-RBC comparison

For experiments designed to directly compare DNA detection in plasma and RBC fractions from cancer patient samples, whole blood was centrifuged at 3,000×g for 10 min. Plasma was carefully removed and aliquoted into appropriately labeled tubes, with 5 µL of plasma transferred into 100 µL DNA/RNA Shield. Care was taken to avoid disturbing the buffy coat and pRBC layer. The remaining plasma and buffy coat were subsequently removed to minimize leukocyte and plasma contamination of the RBC fraction.

For each sample, 5 µL of pRBCs were transferred into 100 µL MACS buffer and mixed thoroughly. Aliquots were centrifuged at 10,000 rpm for 5 min, and the supernatant was carefully removed without disturbing the RBC pellet. The pellet was resuspended in 100 µL MACS buffer and centrifuged again at 10,000 rpm for 5 min. Following removal of the supernatant, the washed RBC pellet was resuspended in 100 µL Zymo buffer and stored at −80°C until further processing.

#### Reticulocyte depletion and isolation of mature RBCs

To evaluate the contribution of reticulocytes to RBC-associated DNA detection, reticulocytes were depleted from the pRBC fraction using anti-CD71 magnetic beads. Briefly, 30 µL of pRBCs was resuspended in 80 µL MACS buffer, followed by addition of 20 µL anti-CD71 magnetic beads (BioLegend). Samples were mixed thoroughly by pipetting and incubated for 15 min at 4°C on a nutator. Following incubation, cells were washed with 1 mL MACS buffer and centrifuged at 300×g for 5 min. The supernatant was removed, and the cell pellet was resuspended in 500 µL MACS buffer.

Magnetic separation was performed using LD Columns (Miltenyi Biotec) positioned in a magnetic separator. Columns were equilibrated with 2 mL MACS buffer before sample application. The labeled cell suspension was applied to the column, and the flow-through containing reticulocyte-depleted cells was collected into a 15-mL conical tube. Columns were subsequently washed twice with 1 mL MACS buffer, and the wash fractions were collected with the flow-through. Cells were centrifuged at 800×g for 5 min and resuspended in 100 µL MACS buffer. Mature RBCs were subsequently isolated using anti-CD235a (glycophorin A) magnetic beads and LS Columns as described above.

#### Isolation of RBCs from mouse whole blood

For mouse experiments, whole blood was collected by cardiac puncture and centrifuged at 1,500×g for 10 min. Plasma was carefully collected and aliquoted into 5-µL fractions, with each aliquot transferred into 100 µL DNA/RNA Shield. The remaining plasma and buffy coat were carefully removed, and the pRBC fraction was retained for RBC isolation.

For magnetic purification, 30 µL of mouse pRBCs were resuspended in 100 µL MACS buffer, and 15 µL anti-TER-119 microbeads (Miltenyi Biotec) were added. Samples were mixed thoroughly by pipetting and processed using the same magnetic labeling, washing, and column separation procedure described above for human RBC isolation, with the appropriate magnetic column and buffer volumes used for the mouse samples. Purified mouse RBCs were subsequently processed according to the requirements of each experiment.

#### Co-culture Experiments

For all co-culture experiments, RBCs isolated from healthy human donors or mice were co-incubated with the human tumor cell lines H1975, H358, and A549 expressing the *EGFR* L858R, *KRAS* G12C, and *KRAS* G12S mutations, respectively. A549 cells were maintained in F-12K medium (Gibco) supplemented with 10% fetal bovine serum (FBS) (Life technologies), whereas H358 and H1975 cells were cultured in RPMI-1640 medium (Gibco) supplemented with 10% FBS. Culture media were replaced every 2–3 days, and cells were passaged upon reaching 70– 80% confluence. All cell lines and base media were obtained from the American Type Culture Collection (ATCC). For co-culture assays, tumor cells (10⁶ cells per well) were seeded in six-well plates and allowed to adhere and reach confluence over 24 h prior to the addition of RBCs.

#### DNA extraction from plasma and RBCs

Genomic DNA was extracted from plasma and RBC samples using the DNeasy Blood & Tissue Kit (Qiagen) according to the manufacturer’s instructions, with modifications to accommodate the sample type and volume. For *in vitro* experiments, DNA was extracted from RBC-containing samples and eluted in 100 µL elution (AE) buffer without further concentration or dilution. For mouse experiments, DNA was extracted from 5 µL of plasma and from a single aliquot containing 10⁷ RBCs. DNA was subsequently concentrated and resuspended in 50 µL AE buffer. For patient samples, DNA was extracted either from pRBCs, or independently from two aliquots of 10⁷ RBCs. Each 10⁷ aliquot was processed separately and eluted in 100 µL AE buffer. The two eluates were then pooled (200 µL total), concentrated, and resuspended in 50 µL AE buffer. Because RBCs contain abundant hemoglobin and heme, residual heme and other RBC-associated components may inhibit DNA amplification and reduce the sensitivity of PCR-based assays (Supp. Fig. 2B). To minimize the potential effects of PCR inhibitors in RBC-derived DNA preparations, serial dilutions of extracted DNA (neat, 1:5, 1:10, and 1:20) were prepared and evaluated by qPCR, as indicated. For qPCR, the dilution producing the optimal amplification performance and cycle threshold (Ct) was selected for subsequent analyses.

#### DNA concentration

DNA samples were concentrated by sodium acetate/ethanol precipitation. Briefly, 1/10 volume of 3 M sodium acetate (pH 5.2) (Thermo Scientific) was added to each DNA sample, followed by vortexing and brief centrifugation. Two volumes of ice-cold 100% ethanol were then added, and samples were vortexed and centrifuged. Samples were incubated at −20°C for 1 h, followed by incubation at −80°C for 30 min. Samples were subsequently centrifuged at 4°C for 30 min at maximum microcentrifuge speed. The supernatant was carefully removed by inverting the tubes over a double layer of paper towels, with sufficient spacing between samples to minimize the risk of cross-contamination. Tubes were left inverted to air-dry for 3 min, taking care not to over-dry the DNA pellet. DNA was then resuspended in 50 µL AE buffer by vigorous vortexing, followed by centrifugation to collect the sample. The resulting 50 µL represented the final concentrated DNA preparation.

#### Quantitative PCR (qPCR)

qPCR was performed using TaqMan 2× Universal Master Mix (Applied Biosystems) on a QuantStudio 6 Flex Real-Time PCR System (Applied Biosystems) using previously established cycling conditions. A549, H358, and H1975 tumor cell lysates were used as positive controls for the corresponding assays. Appropriate no-template controls were included in each experiment. For *in vitro* experiments, 100 µL of extracted DNA were used directly as the qPCR template without further dilution or concentration. For mouse and patient samples, the extracted DNA (±concentration) was analyzed using the dilution series described above to minimize the effects of potential RBC-associated PCR inhibitors. The dilution yielding the optimal Ct and amplification performance was selected for downstream analysis.

For absolute copy-number quantification by qPCR, tumor cells (10⁶) were plated overnight, subsequently trypsinized, and processed for genomic DNA extraction. The target genomic region was amplified by conventional PCR, and the resulting amplicon was separated by agarose gel electrophoresis. The target band was excised and purified using a gel extraction kit (Thermo Fisher), and DNA concentration was quantified using a NanoDrop spectrophotometer. The purified amplicon was used to generate a standard curve consisting of six serial 10-fold dilutions of known DNA concentration (Supp. Fig. 1C). Standards were run in parallel with experimental samples on the same qPCR plate. Absolute target copy number was determined by comparing the Ct values of experimental samples with the standard curve generated from the known amplicon standards. Final copy-number calculations accounted for amplicon size, DNA input volume, and applicable dilution factors. qPCR data for ES patients were previously reported(*5*), but extraction and qPCR/ddPCR were repeated here with our newly optimized RBC method, and updated data are presented in this manuscript.

#### Droplet digital PCR (ddPCR)

ddPCR was performed using the Bio-Rad digital PCR platform with the appropriate forward and reverse primers and fluorescently labeled probes. DNA templates consisted of concentrated RBC-derived DNA corresponding to 2×10⁷ RBCs, or 5 µL of plasma-derived DNA vs 5 µL of RBC-derived DNA, as indicated for each experiment. For each reaction, 8–10 µL of DNA template was combined with the appropriate ddPCR master mix and primer/probe set. Reaction mixtures were partitioned into nanoliter-sized droplets and subjected to PCR amplification using the Bio-Rad ddPCR system according to the manufacturer’s recommended cycling conditions. Following amplification, droplets were analyzed using a Bio-Rad droplet reader. Fluorescence signals were used to distinguish target-positive and wild-type droplets, and target concentrations were calculated using Bio-Rad analysis software with Poisson correction. ddPCR quality control criteria were met for all samples, with >20,000 accepted droplets generated per reaction. The reported absolute target copy numbers were calculated by multiplying the concentration by the relevant final sample volume. Cancer cell lysate-derived DNA was used as a positive control, and no-template controls were included in each ddPCR experiment.

#### Transwell Assays

Tumor cells (10⁶) were seeded and cultured for 24 h. Media were subsequently removed, and cells were washed with PBS. RBCs (10⁷) were then co-cultured either directly with tumor cells or indirectly using 24 mm Transwell inserts with 0.4 μm polycarbonate membranes (Corning). Following 24 h of co-culture, RBCs were collected, washed with PBS, and subjected to DNA extraction. qPCR was performed to assess the detection of the tumor-specific mutations.

#### Stability of rtDNA

Tumor cells (10⁶) were seeded and cultured for 24 h. The media were then removed, and cells were washed with PBS. Subsequently, RBCs (10⁷) were added and co-cultured with tumor cells for an additional 24 h. Following co-culture, RBCs were collected and washed with PBS, then processed immediately for DNA extraction. qPCR was performed weekly on DNA extracted prior to storage to assess the detection and stability of the tumor-specific mutations.

#### Minimum RBC Input Required for rtDNA Detection

Tumor cells (10⁶) were seeded and cultured for 24 h. The media were then removed, and cells were washed with PBS. RBCs were serially diluted to concentrations of 10⁷, 10⁶, 10⁵, 10⁴, and 10³ cells and co-cultured with tumor cells for 24 h. Following co-culture, RBCs were collected, washed with PBS, and subjected to DNA extraction. Mutation-specific qPCR was then performed to quantify tumor-derived DNA.

#### DNA Acquisition Kinetics Experiments

Tumor cells (10⁶) were seeded, cultured for 24 h, washed with PBS, and co-cultured with RBCs (10⁷) for varying durations (5 min, 15 min, 30 min, 1 h, 2 h, and 4 h). RBCs were collected at each time point, washed with PBS, and processed for DNA extraction. Mutation-specific qPCR analysis was performed to evaluate the temporal dynamics of DNA acquisition.

#### DNase Treatment Assays

Tumor cells (10⁶) were seeded and cultured for 24 h, washed with PBS, and co-cultured with RBCs (10⁷, 10⁶, 10⁵, or 10⁴) for 24 h. Following co-culture, RBCs were collected, washed with PBS, and divided into two equal fractions. Cell pellets were resuspended either in PBS containing Ca²⁺ and Mg²⁺ (no DNase control) or in PBS supplemented with 500 μg/mL DNase I (Millipore Sigma). Samples were incubated for 2 h at 37°C on a nutator. After incubation, cells were pelleted, washed twice with PBS, and subjected to DNA extraction. Mutation-specific qPCR was performed to assess the impact of DNase treatment.

#### TLR9 inhibition/Neuraminidase (NA) treatment

Tumor cells (10⁶) were seeded and cultured for 24 h, then washed with PBS. RBCs were isolated as above, and the resulting pellets were resuspended in media supplemented with 0.05 µg IgG, 0.05 µg anti-TLR9 antibody (clone 5G5; Abcam), or 0.5 U/mL NA (Sigma Aldrich) in combination with either IgG (Biolegend) or anti-TLR9 antibody. Samples were incubated for 2 h at 37°C on a nutator. Following incubation, cells were pelleted, washed twice with PBS, and co-cultured with tumor cells for an additional 24 h. Following co-culture, RBCs were collected, washed with PBS, and subjected to DNA extraction. Mutation-specific qPCR was performed.

#### Flow cytometry

RBCs were isolated and washed with PBS or Fluorescence-Activated Cell Sorting (FACS) buffer (Rockland Immunochemicals Inc.) prior to use. Where indicated, RBCs (10⁶) were treated with NA (0.5 U/mL) in PBS for 2 h at 37°C, followed by a single wash with FACS buffer (800×g, 5 min). For co-culture experiments, tumor cells (10⁶) were seeded and cultured for 24 h, washed with PBS, and then co-cultured with RBCs (10⁶) for an additional 24 h. RBCs were incubated with Sytox (1 µM) (Invitrogen) or FACS buffer alone for 30 min in the dark at room temperature under gentle agitation, then washed twice with FACS buffer. For CpG stimulation assays, NA-treated or untreated RBCs were resuspended in PBS and incubated with HPV-CpG-ATTO-labeled oligonucleotide (0.5 μM or 2 µM) (Integrated DNA technologies) for 2 h at 37°C, followed by a single wash with FACS buffer. For detection of tumor DNA on cancer cell surface, tumor cells (10⁶) were seeded overnight, stained with Sytox (1 µM) for 30 min, incubated with DNAse or RBCs for 2 h, then processed for flow imaging. For surface staining, cells were incubated with anti-TLR9-FITC (clone 5G5; 1 µg) or an isotype-matched control (IgG2a-FITC; 1µg) at room temperature for 1 h under gentle agitation. All samples were pelleted by centrifugation, washed as indicated, and analyzed on the flow cytometer (LSR Fortessa or Cytoflex). Data were analyzed using FlowJo (v10.10.1).

#### Murine Models of lung cancer

All animal procedures were conducted in compliance with Institutional Animal Care and Use Committee (IACUC) guidelines at the University of Pennsylvania (Animal Welfare Assurance protocol #804000). Female NOD/scid/IL2rg (NSG) mice were bred in the Penn Stem Cell and Xenograft Core and were used for experiments at 10 to 16 weeks of age. The A549-NY-ESO-luc cell line was generated by transducing the human cell line A549 with lentiviral plasmid containing NY-ESO, HLA-A2, and luciferase(*17*). For the intravenous injection group, A549-NY-ESO-luc cells (10⁶) were administered via the tail vein. Tumor growth was monitored by bioluminescence imaging using an IVIS Spectrum imaging system (PerkinElmer). Bioluminescent signals were acquired with automatic exposure settings and quantified as total photon flux (photons/sec) within a defined region of interest (ROI) using Living Image software (PerkinElmer). Background signal was subtracted using ROIs from non-tumor-bearing regions. For the subcutaneous injection group, A549-NY-ESO-luc cells (10⁶) were injected into the flanks of NSG mice. Peripheral blood (30– 50 µL) was collected from the lateral tail vein of each animal into EDTA-coated microcentrifuge tubes on all time points. Animals were then euthanized, and terminal blood samples (100-200 µL) were obtained via cardiac puncture for the final time point.

#### Patient Recruitment and Sample Preparation

Patients presenting to the thoracic oncology clinic at the Hospital of the University of Pennsylvania with NSCLC provided written informed consent for blood collection and use of clinical data. 10 mL of whole blood was collected, and RBCs were isolated within 2 hours of sample receipt. Details of the patient cohorts are provided in Tables 1 and 2. DNA was extracted from RBCs and plasma as described in this Methods section above. Patient oncologic characteristics, including somatic mutation status, were obtained by reviewing the electronic medical record. This study was approved by the Institutional Review Board at the University of Pennsylvania (IRB# 855824).

### Statistical analysis

Differences between groups were compared using a t-test, one-way analysis of variance (ANOVA), and two-way ANOVA as appropriate. Multiple comparisons tests including Tukey’s, Šidák’s, Wilcoxon, and Mann-Whitney were used for post hoc analysis. All statistical analyses were performed using GraphPad Prism version 11. P value <0.05 was considered significant for all analyses.

### Data Preparation and Language Editing

All subfigures were generated using GraphPad Prism version 11 and assembled into final figures using Adobe Illustrator version 30.8.2. Flow data were generated in the Penn Cytomics and Cell Sorting Shared Resource Laboratory at the University of Pennsylvania (RRID:SCR_022376). Penn Cytomics is partially supported by the Abramson Cancer Center NCI Grant (P30 CA016520). Artificial intelligence (AI) tools were used solely for language editing to improve clarity, grammar, and readability. No AI tools were used to generate, analyze, or interpret scientific data or content.

## Notes

### Summary of Updates

An error in one of the references has been updated.

