## Supplementary Figures for "The Red Blood Cell Surface Defines a Distinct Tumor DNA Compartment for Liquid Biopsy"

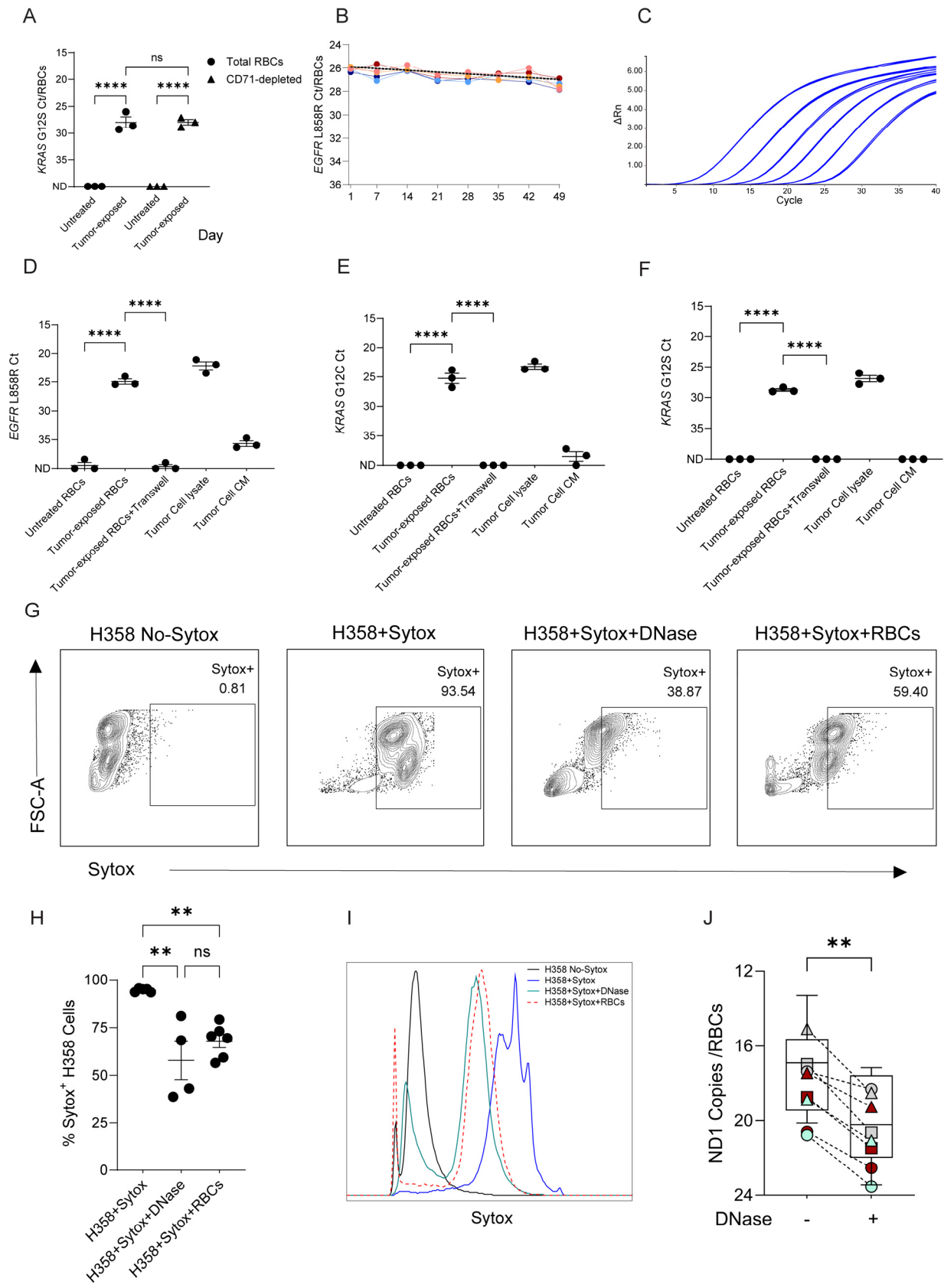

**Figure S1. rtDNA is stable, acquired through close proximity, and associates with the surface of RBCs.** (A) Detection of tumor-derived DNA in total and reticulocyte-depleted RBCs (CD71-depleted) following co-culture with A549 cells, showing no significant difference between populations. n=3 HDs, 2 independent experiments. Statistical analysis performed with one-way ANOVA with Šidák's multiple comparisons test. ND: Not Detected. (B) RBCs from HDs were co-cultured with H1975 cells. DNA extracted from RBCs was stored at $-20^{\circ}\text{C}$ and analyzed weekly by qPCR for 7 weeks. Individual donor trajectories are shown in color; dashed line represents the mean. The RBC-bound DNA signal remained stable through 42 days (linear regression of group means: slope=0.015 per day, $R^2=0.54$, $P=0.059$) but showed a significant decrease by day 49 (slope=0.022 per day, $R^2=0.69$, $P=0.011$). n=5 HDs. (C) Representative qPCR standard curve. (D-F) qPCR detection of *EGFR* L858R, *KRAS* G12C, and *KRAS* G12S mutations in RBCs following direct co-culture, transwell co-culture, or conditioned media (CM) exposure to H1975, H358, and A549 cells, respectively. n=3 HDs, 3 independent experiments. Statistical analysis performed using one-way ANOVA with Šidák's multiple comparisons. (G) Representative contour plots showing Sytox staining of H358 cells under control conditions and following co-incubation with RBCs or DNase treatment. n=6 HDs, one representative experiment shown. (H) Quantification of Sytox<sup>+</sup> H358 cells following Sytox staining in the presence or absence of RBCs or DNase treatment. n=6 HDs, 3 independent experiments. Data are presented as mean ± SEM. Statistical analysis was performed using one-way ANOVA with Tukey's multiple comparisons test. (I) Representative histograms of Sytox fluorescence corresponding to the conditions shown in G. (J) DNase treatment of RBCs following co-culture with H1975, H358, or A549 cells significantly reduced mtDNA (ND1). Squares represent H1975 co-culture, circles represent H358, and triangles represent A549. Each color represents an individual RBC donor (n=3 HDs), 3 independent experiments. Data are presented as mean ± SEM. Statistical analysis was performed using the Wilcoxon matched-pairs signed-rank test. \*\*$P<0.01$, \*\*\*\*$P<0.0001$.

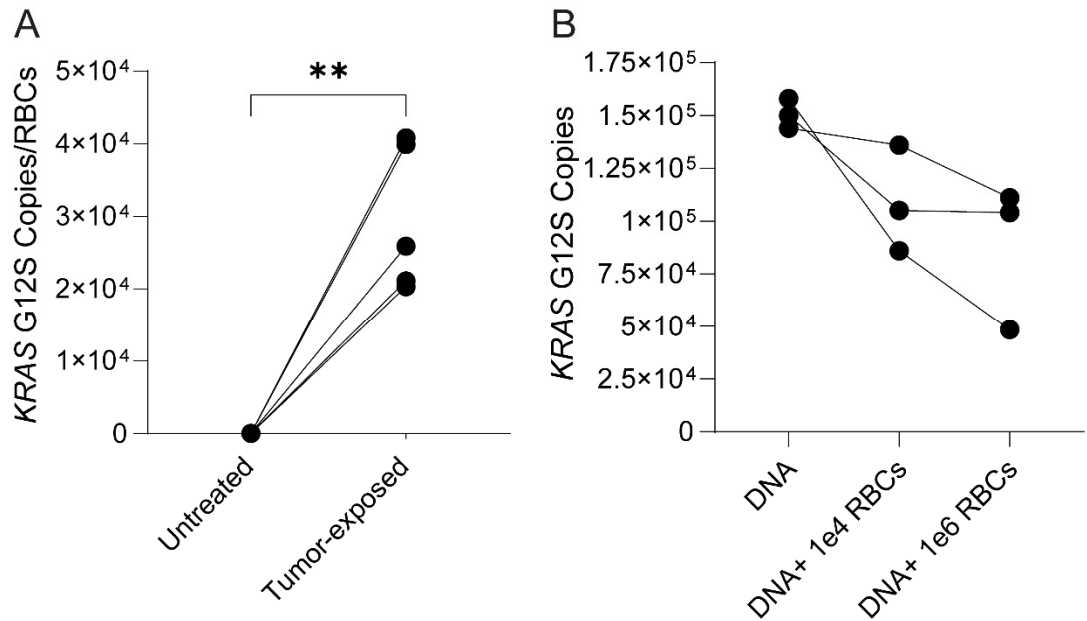

**Figure S2. G12S mutant DNA from A549 cells is detected on mouse RBCs following co-culture, while RBC-associated factors inhibit DNA amplification. (A)** qPCR detection of *KRAS* G12S mutant DNA on murine RBCs following co-culture with A549 cells. n=5 mice, 3 independent experiments. Statistical analysis performed using a paired t-test. **(B)** RBC-associated inhibition of DNA amplification. To investigate whether inhibitors within RBCs can interfere with PCR DNA amplification (potentially by inhibiting DNA polymerase activity and/or interfering with the reaction environment), 100 μL of extracted genomic DNA from A549 cells harboring the *KRAS* G12S mutation was combined with increasing concentrations/dilutions of RBCs. DNA amplification was subsequently assessed to determine whether exposure to RBCs affected detection of the G12S target on cancer cells. Exposure to two different increasing amounts of RBCs resulted in progressively reduced G12S signal, indicating inhibition of DNA amplification in the presence of RBC-associated inhibitory components. This effect may be attributable to RBC-derived factors, including heme and other intracellular components released from RBCs, which can interfere with DNA amplification and/or detection. n=3 HDs. \*\*P<0.01

**Table S1. Sequences and IDs of qPCR and ddPCR primers**

| **qPCR primers (IDT)** | **Sequence** |
| --- | --- |
| G12S Forward | 5'-AATATAAACTTGTGGTAGTTGGAGCgA-3' |
| G12C Forward | 5'-AATATAAACTTGTGGTAGTTGGAGCcT-3' |
| G12R Forward | 5'-AATATAAACTTGTGGTAGTTGGAGCtC-3' |
| KRAS common Reverse | 5'-CATATTCGTCCACAAAATGATTCTG-3' |
| *KRAS* probe | 5'-FAM-CTGTATCGTCAAGGCACT-MGB-3' |
| EGFR Forward | 5'-GCAGCATGTCAAGATCACAGATT-3' |
| EGFR Reverse | 5'-CCTCCTTCTGCATGGTATTCTTTCT-3' |
| L858R Probe | 5'-AGTTTGGCCCGCCCAA-3' |
| ND-1 Forward | 5'-ATACCCATGGCCAACCTCCT-3' |
| ND-1 Reverse | 5'-GGGCCTTTGCGTAGTTGTAT-3' |
| **ddPCR primers (BioRad)** | **ID** |
| EGFR WT | dHsaCP2000022 |
| EGFR L858R | dHsaCP2000021 |
| KRAS G12C | dHsaMDV2510584 |
| KRAS G12R | dHsaMDV2510590 |
